# Use of target-displaying magnetized yeast in screening mRNA display peptide libraries to identify ligands

**DOI:** 10.1101/2020.08.13.249722

**Authors:** John Bowen, Kaitlyn Bacon, Hannah Reese, Balaji M. Rao, Stefano Menegatti

**Author notes:** denotes equal contribution (John Bowen, Kaitlyn Bacon).

## Abstract

This work presents the first use of yeast-displayed protein targets for screening mRNA display libraries of cyclic and linear peptides. The WW domains of Yes-Associated Protein 1 (WW-YAP) and mitochondrial import receptor subunit TOM22 were adopted as protein targets. Yeast cells displaying WW-YAP or TOM22 were magnetized with iron oxide nanoparticles to enable the isolation of target-binding mRNA-peptide fusions. Equilibrium adsorption studies were conducted to estimate the binding affinity (K_D_) of selected WW-YAP-binding peptides: K_D_ values of 37 μM and 4 μM were obtained for cyclo[M-AFRLC-K] and its linear cognate, and 40 μM and 3 μM for cyclo[M-LDFVNHRSRG-K] and its linear cognate, respectively. TOM22-binding peptide cyclo[M-PELNRAI-K] was conjugated to magnetic beads and incubated with yeast cells expressing TOM22 and luciferase. A luciferase-based assay showed a 4.5-fold higher binding of TOM22^+^ yeast compared to control cells. This work demonstrates that integrating mRNA and yeast display accelerates the discovery of biospecific peptides.

---

Cell-free combinatorial libraries, such as ribosomal- and mRNA-display, are attractive tools in ligand discovery owing to their chemical diversity and ease of post-translational modifications.^1^ In the mRNA-display platform, a peptide sequence is connected to its coding mRNA via a puromycin linkage.^2^ The mRNA is reverse transcribed to form mRNA-cDNA-peptide fusions that can be used in library screening to identify peptide affinity ligands. Following library selection, the DNA linked to the candidate peptide binders is amplified and sequenced. This technology is widely utilized in the discovery of short peptides,^3^ which are of great interest for use as tags or modulators of protein-protein interactions;^4^ short peptides are also amenable to post-translational modifications, such as cyclization, which typically imparts higher target binding affinity and proteolysis resistance.^5^ In prior work, our group developed a method to generate mRNA-display libraries of cyclic peptides via head-to-side chain chemical crosslinking of peptide-mRNA fusions adsorbed on a solid phase.^6^

The selection of mRNA-display peptide libraries is commonly performed against a target protein conjugated onto synthetic magnetic nanoparticles.^6,7^ In other screening techniques, such as phage- and yeast-display, the use yeast displayed targets is becoming prevalent.^8^ Target expression on the surface of yeast cells eliminates the need to separately produce and isolate the target protein and is particularly impactful when considering species prone to misfolding when recombinantly produced in hosts like *E.coli*.^9,10^ For example, the extracellular domain of membrane proteins, have been expressed using the yeast display platform in the context of structure-function studies and for the screening of phage- and yeast-display libraries.^8^

Integrating yeast-displayed targets with the screening of mRNA-display libraries, however, poses a challenge: how to separate the selected fusions from the unbound portion of the library. As the fusions are soluble, the use of centrifugation for separating yeast-bound fusions represents an option; however, the carryover of unbound fusions during centrifugation is consistently observed, resulting in a strong contamination of false positives. This highlights the need for improved isolation techniques capable of reducing carryover and increasing selection stringency.

To address this challenge, this work presents the use of magnetized yeast cells as protein-display particles to increase the efficiency of isolating lead mRNA-peptide fusions (**Figure 1**). Yeast are magnetized by adsorbing iron oxide nanoparticles (IONs) onto anionic moieties that constellate on the cell wall in a pH-controlled environment.^11^ Yeast expressing proteins with varying isoelectric points can be magnetized by adjusting the pH of the buffer in which the IONs are adsorbed.^12^ Following adsorption, the IONs are blocked with albumin to prevent non-specific adsorption of mRNA-peptide hybrids and other species in solution. The selection of mRNA-display libraries generally takes place at a pH of 7.4. If yeast are magnetized at a different pH, IONs can dissociate when the magnetic yeast cells are incubated in the selection buffer, and all mRNA-peptide hybrids associated with a demagnetized yeast cell are lost. It has nonetheless been shown more than 50% of non-specifically magnetized cells maintain their magnetized state over the course of a selection.^8^ The size of mRNA-display libraries (~10^12^ hybrids, corresponding to ~1000 copies per peptide in case of 7-mers) and the number of target proteins expressed on yeast (~50,000 per cell) ensure ample contact between the library and the target proteins. Furthermore, because there are multiple copies of each peptide in the library, it is likely that any peptide with affinity for the target can be isolated even if some fusions are lost due to ION desorption.

**Figure 1.**
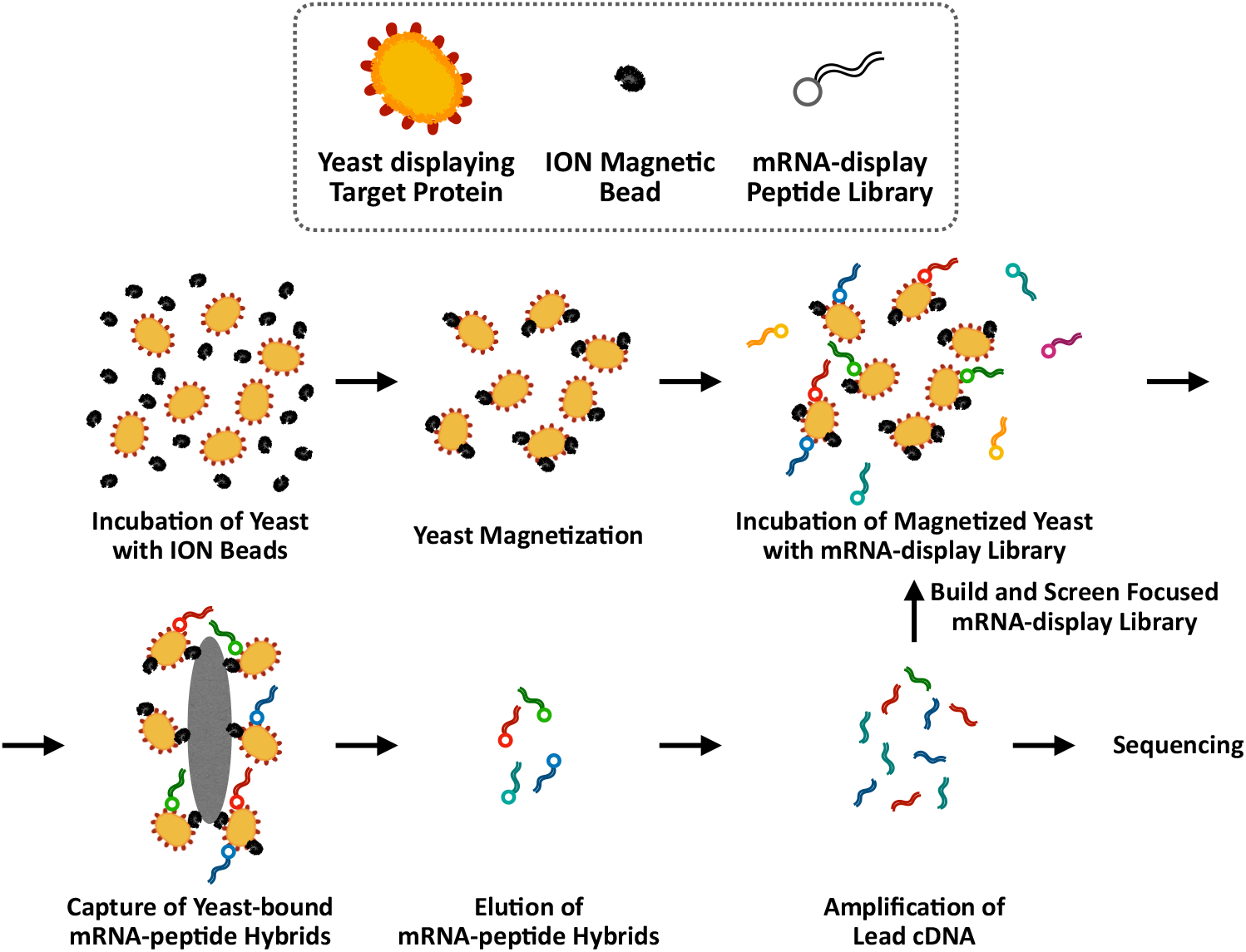
Non-specific yeast cell magnetization and mRNA-display-peptide library screening against magnetic yeast targets. Yeast cells expressing WW-YAP or TOM22 were incubated with iron oxide magnetic nanoparticles (ION magnetic beads). Magnetic yeast cells were separated from non-magnetized yeast cells using a magnet. The mRNA-display-peptide library was then incubated with the magnetized target cells. The mRNA-display peptide fusions bound to the target cells were isolated using a magnet. The positively bound mRNA-display peptide fusions were eluted from the target cells. DNA linked to the eluted clones was amplified, and a new library was synthesized using this DNA as the template for the next round of screening. The selection process was repeated multiple rounds against the target cells. The stringency of elution increased for each selection round.

In this work, a combinatorial library of cyclic peptides generated using mRNA-display was screened against magnetized target yeast to identify affinity peptides to the tandem WW domain of Yes-Associated Protein 1 (WW-YAP) and mitochondrial surface protein TOM22. YAP is an effector protein of the Hippo Signaling pathway regulating cellular amplification and survival^13^, while TOM22 is a mitochondrial membrane protein involved in the recognition and translocation of mitochondrial pre-proteins produced in the cytosol.^14^ Peptides capable of inhibiting the binding of cytoplasmic proteins to the WW domains of YAP can provide mechanistic insight into the Hippo-YAP pathway and a means to regulate cell proliferation. Peptides with affinity for TOM22 can be used for the targeted delivery of therapeutic payloads to treat patients with mitochondrial deficiencies.^15^ WW-YAP and TOM22 were expressed on the surface of EBY100 yeast using the yeast surface display platform.^16^ The cells were magnetized with IONs and utilized as targets during subsequent selections of mRNA-display peptide libraries.

A mRNA-display library of cyclic peptides was constructed following a previously developed method of head-to-tail cyclization that employs a glutarate linker to tether amine groups within methionine and lysine residues flanking the variable peptide segment.^6,17^ Prior to selecting against target yeast cells, the library was negatively screened against magnetized yeast displaying a non-target protein (*i.e*., Fc portion of human IgG, hFc) to remove fusions that bound to other proteins on the yeast surface. Throughout the library selection process, the stringency of the elution conditions was increased to promote isolation of high-affinity binders. Initially, the mRNA-peptide fusions were eluted from the target cells using 0.15 M potassium hydroxide; in later rounds, the yeast-bound peptides were washed with a mild acidic buffer (50 mM NaCl in 50 mM sodium acetate at pH 5) prior to alkaline elution to remove weakly bound peptides and increase the probability of isolating true affinity binders. Following elution, the cDNA linked to the isolated mRNA-peptide fusions was amplified to generate a focused sub-library for the subsequent screening rounds. We noted that, during the generation of the sub-libraries, heptamer and decamer sequences emerged. While potentially present as minor contaminants in the original library, these longer peptides outcompeted the original pentamers, owing to their higher length-driven enthalpic contribution to the binding energy, and were identified as the lead fusions after multiple rounds of selection.

Sequencing returned 10 candidate WW-binding peptides: 3 pentamers, 2 heptamers, and 5 decamers (**Figure 2A**). Two pentamer sequences were discarded due to a lack of chemical diversity (*i.e*., predominance of aliphatic residues combined with a lack of ionic, hydrogen-binding, and aromatic residues). Both heptameric sequences were discarded, one for containing cysteine, which can lead to covalent peptide-protein disulfide bonds, and the other due to an excess of aliphatic residues. Of the decameric sequences, two were undetermined (*i.e*., Sanger sequencing returned unassigned residues “X”), and one featured poor chemical diversity (*i.e*., predominance of aliphatic residues).

**Figure 2.**
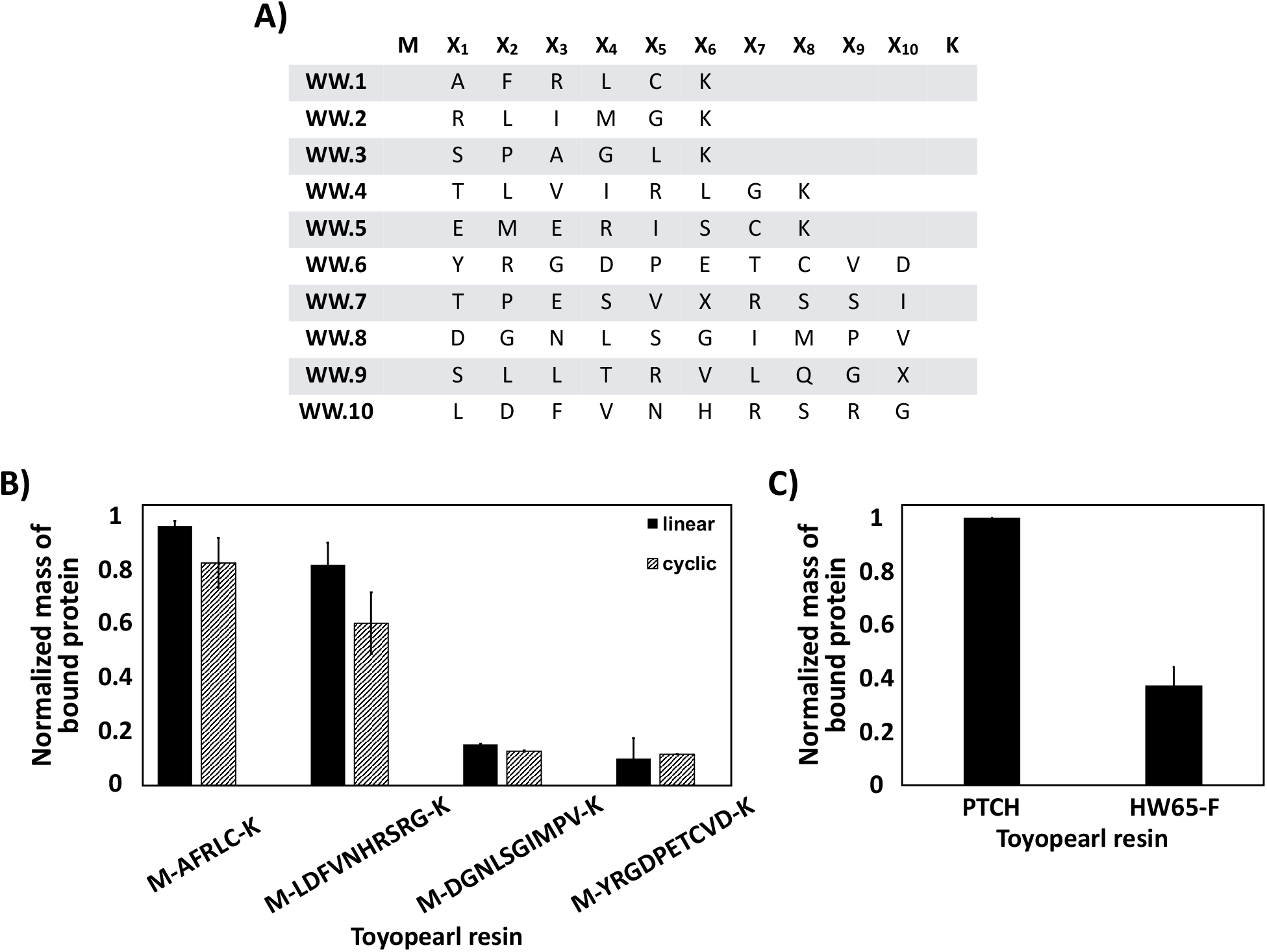
Identified WW-YAP-binding cyclic peptides and primary evaluation of lead candidates. **(A)** Amino acid sequences of peptides isolated from screening the mRNA-display library of cyclic peptides against yeast cells displaying WW-YAP. (**B**) Soluble WW-YAP protein (1 mg/mL) was incubated with cyclo[M-AFRLC-K]-Toyopearl, MAFRLCK-Toyopearl, cyclo[M-LDFVNHRSRG-K]-Toyopearl, MLDFVNHRSRGK-Toyopearl, cyclo[M-DGNLSGIMPV-K]-Toyopearl, MDGNLSGIMPVK-Toyopearl, cyclo[M-YRGDPETCVD-K]-Toyopearl, and MYRGDPETCVDK-Toyopearl resins; the amount of unbound protein in solution was determined by measuring the A280 absorbance of the supernatant, and the amount of protein bound by each resin was calculated via mass balance. **(C)** RYSPPPPYSSHS-Toyopearl (PTCH peptide, positive control) and Toyopearl HW65-F (negative control) resins were also incubated with WW-YAP protein (1 mg/mL). In panels (**B**) and (**C**), the mass of WW-YAP protein bound by each resin was normalized to the mass of WW-YAP protein bound by RYSPPPPYSSHS-Toyopearl resin. Error bars correspond to the standard error of the mean from three independent replicates.

We therefore conducted a primary evaluation of four cyclic peptides, cyclo[M-AFRLC-K], cyclo[M-LDFVNHRSRG-K], cyclo[M-DGNLSGIMPV-K], and cyclo[M-YRGDPETCVD-K], via equilibrium binding tests using recombinant human WW-YAP (1 mg/mL). The cyclic peptides and their linear cognates were synthesized on solid phase (Toyopearl amino AF-650M) via standard Fmoc/tBu chemistry; Toyopearl resin functionalized with the known WW-binding peptide PTCH (RYSPPPPYSSHS)^18,19^ and hydroxyl Toyopearl HW-65F resin were adopted as positive and negative control adsorbents, respectively. Sequences AFRLC and LDFVNHRSRG showed significant WW-YAP binding, capturing 60% and 44% of the protein in solution in their cyclic form, and 68% and 60% in their linear form, respectively (**Figure 2B**); similarly, PTCH-Toyopearl resin bound > 73% of the incubated WW-YAP (**Figure 2C**). In contrast, sequences DGN-LSGIMPV and YRGDPETCVD showed poor WW-YAP binding and were discarded.

Selected sequences AFRLC and LDFVNHRSRG were then characterized via adsorption isotherm studies to evaluate their binding strength and capacity. The assays were performed by incubating WW-YAP at varying concentrations (0.02 – 12.5 mg/mL) with PTCH-Toyopearl, MAFRCLK-Toyopearl, cyclo[M-AFRLC-K]-Toyopearl, MLDFVNHRSRGK-Toyopearl, and cyclo[M-LDFVNHRSRG-K]-Toyopearl resins. The values of adsorbed protein (*Q*, mass of WW-YAP bound per mL of resin) *vs*. equilibrium concentration in solution (*C\**, mass of WW-YAP per mL) were fit against model isotherm equations. The isotherms and the corresponding values of binding capacity (*Q_max_*) and affinity (K_D_) are presented in **Figure 3**. In all cases, a Langmuir-Freundlich behavior was observed, wherein a Langmuir-only profile is observed at *C\** < 2 mg/mL, and the Freundlich profile becomes dominant at *C\** > 2 mg/mL (**Figure S1**). It was noted that the linear form of the selected sequences possess a higher affinity for WW-YAP than their corresponding cyclic form; while cyclic peptides generally feature higher affinity compared to their linear counterparts, instances have been reported of linear peptides with higher protein-targeting activity.^20^

**Figure 3.**
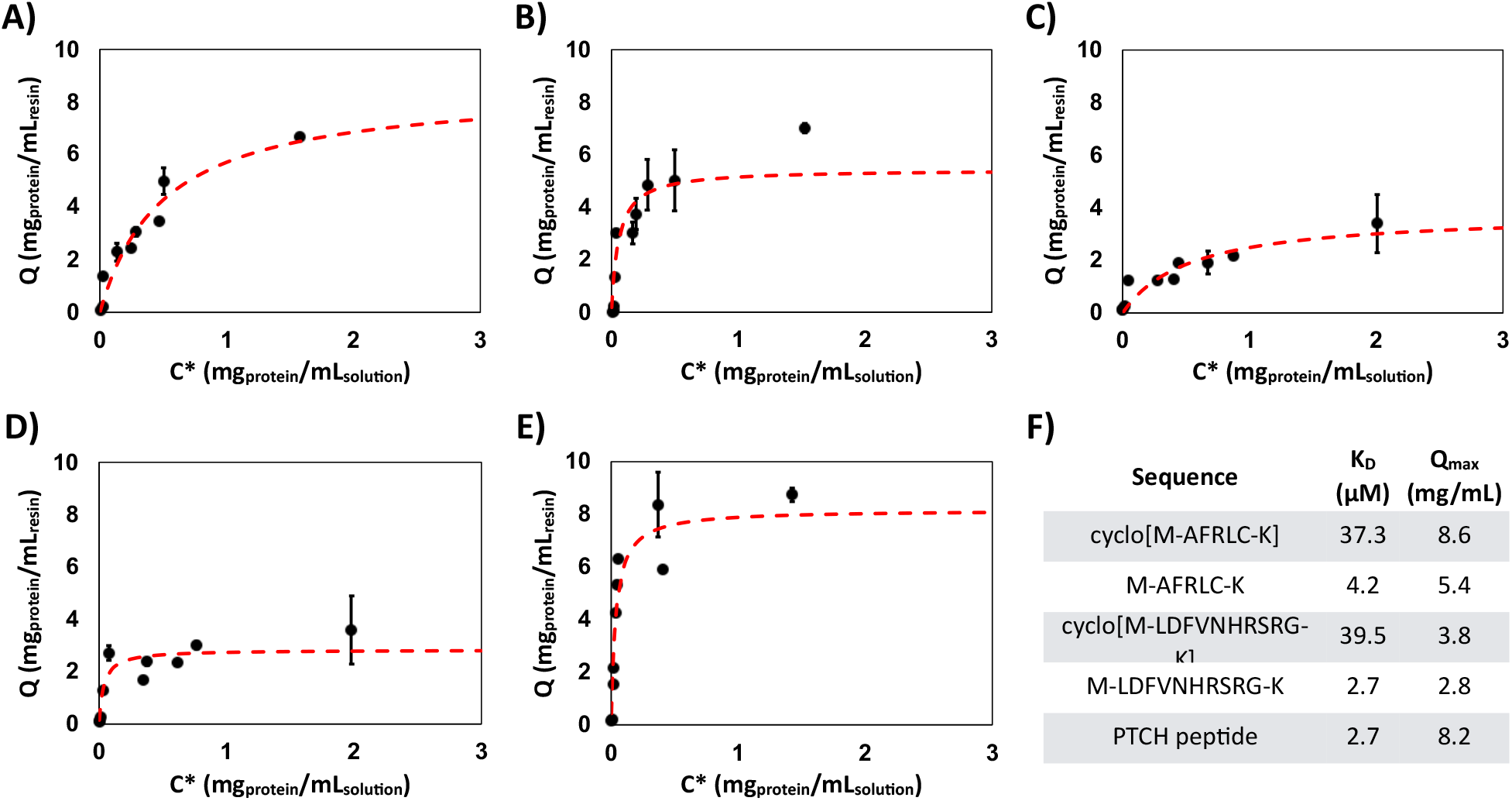
WW-YAP Equilibrium binding isotherms for (**A**) cyclo[M-AFRLC-K]-Toyopearl, (**B**) MAFRCLK-Toyopearl, (**C**) cyclo[M-LDFVNHRSRG-K]-Toyopearl, (**D**) MLDFVNHRSRGK-Toyopearl, and (**E**) RYSPPPPYSSHS-Toyopearl resins. The binding isotherms were generated by plotting the amount of protein bound to the resin (*Q*) *vs*. the concentration of protein in solution at equilibrium (*C\**). **(E)** Resulting values of affinity (K_D_) and maximum protein binding capacity (*Q_max_*) derived from fitting the adsorption data corresponding to *C\**,ranging from 20 μg/mL to 2 mg/mL of WW-YAP, against a Langmuir isotherm equation.

Asymptotic Langmuir behavior plateauing above *C\** of 2 mg/mL was replaced by linear increasing Freundlich behavior (**Figure S1**), which can be imputed to the aggregation of WW-YAP proteins. WW domains are structured as a bundle of 3 antiparallel β-sheets.^21–23^ As the concentration in solution increases, free WW-YAP can self-assemble onto the initial cohort of peptide-bound WW, thereby promoting the formation of an aggregated layer. As the WW-YAP:WW-YAP interaction is likely weaker than the WW-YAP:peptide affinity bond, the binding isotherms exhibit a Freundlich profile at higher concentrations, where significant WW-YAP adsorption occurs by aggregation. **Figure S1** showcases the binding isotherms combining Langmuir and Freundlich profiles, while **Table S1** reports the resulting binding parameters, including binding affinity (K_D_), maximum binding capacity (*Q_max_*), distribution coefficient (K), and correction factor (n).

Finally, the use of cyclo[M-AFRLC-K]-Toyopearl and cyclo[M-LDFVNHRSRG-K]-Toyopearl resins as adsorbents for protein purification in bind-and-elute mode was evaluated (**Table S2**). Notably, complete recovery of bound WW-YAP was achieved using a low-pH glycine buffer. This demonstrates the utility of these identified cyclic peptides as potential purification ligands.

We also investigated the binding strength and orientation of peptides MAFRCLK-GSG, cyclo[M-AFRLC-K]-GSG, MLDFVNHRSRGK-GSG, cyclo[M-LDFVNHRSRG-K]-GSG, and the PTCH peptide (RYSPPPPYSSHS) *in silico* via molecular docking. The crystal structures of the peptides, initially generated by molecular dynamics (MD), were docked against the WW-YAP (PDB ID: 2LTW) using HADDOCK (High Ambiguity Driven Protein-Protein Docking, v.2.2).^24^ The GSG (Gly-Ser-Gly) spacer located on the C-terminus of the peptides was marked as “inactive” to ensure its outward orientation in the WW:peptide complexes; the GSG spacer is utilized to link the peptides on the Toyopearl resin and is not involved in binding. The resulting clusters for each WW:peptide complex were ranked using the scoring functions FireDock and XScore^25^ and the top binding poses were subsequently evaluated by atomistic MD simulations to evaluate the free energy of binding (ΔG_b_). Representative examples of modeled WW:peptide complexes are reported, and the values of K_D_ calculated from the averaged values of binding energy (ΔG_B_) are listed **(Figure 4)**. The docking results indicate that all selected peptides share the WW binding domain of the Smad7 (GESPPPPYSRYPMD, PDB ID: 2LTW)^26^ and PTCH peptides^18^, both of which feature the WW-binding PPXY motif (X: any guest amino acid). The calculated values of K_D_ are in good agreement with those determined through binding isotherms, especially in that the linear peptides bind WW-YAP stronger than their cyclic counterparts. The slightly higher affinity predicted by the MD simulations compared to the results of the binding isotherms can be attributed to the nonideality of the binding events on the chromatographic resin. *In silico*, the WW-YAP target and the peptide ligand enjoy complete rotational freedom and lack of topological constraints, except for defining GSG as a passive linker. In contrast, the binding on the chromatographic resin is influenced by the roughness of the surface and the orientation of the peptides conjugated on it.

**Figure 4.**
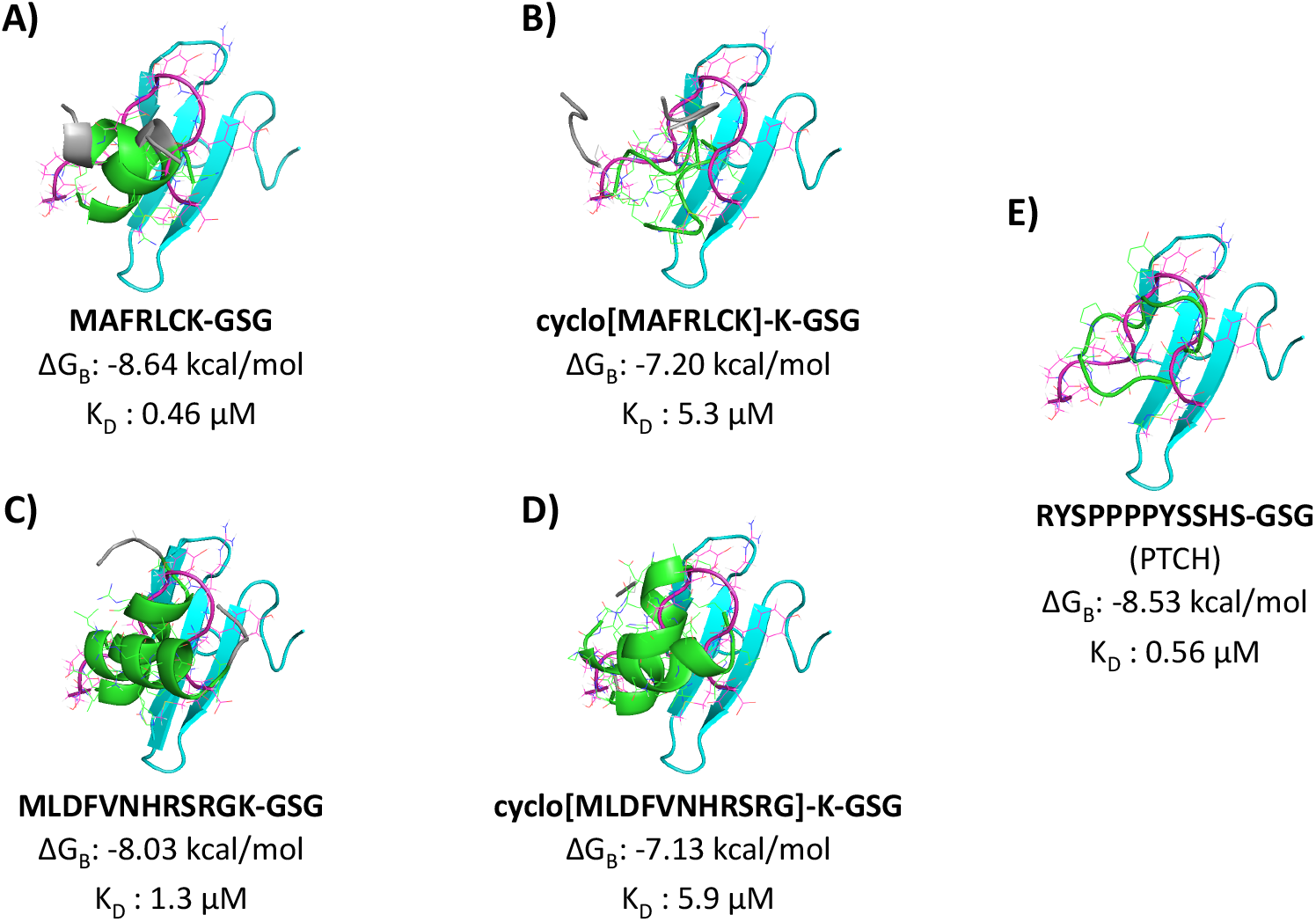
*In silico* WW:peptide complexes obtained by molecular docking and refined by molecular dynamics simulations. **(A)** MAFRLCK-GSG, **(B)** MLDFVNHRSRGK, **(C)** cyclo[M-AFRLC-K]-GSG, **(D)** cyclo[M-LDFVNHRSRG-K]-GSG, and **(E)** PTCH peptide. WW-YAP is denoted in cyan, while the peptides discovered in this work are designated in green with their GSG segment in grey and the Smad7 peptide in magenta. The values of binding energy (ΔG_B_) and corresponding affinity (K_D_) of the WW:peptide complexes derived from the docking and MD simulations are noted. The binding energy and affinity estimates for the WW-PTCH peptide as predicted by molecular simulation are similar to values previously predicted using Isothermal Titration Calorimetry, −6.24 kcal/mol and 27.6 μM, respectively.^18^ The interaction of another WW binding peptide, SMAD7, was also evaluated using molecular docking and MD simulation (*in silico* complex not shown). The binding energy and binding affinity of the WW:SMAD7 complex was estimated as −8.2 kcal/mol and 0.99 μM, respectively.

The mRNA-display selection process was repeated to identify affinity peptides against TOM22. After five rounds of selection, seven sequences were identified (**Figure 5A**). The majority of these sequences were enriched in cationic amino acids, which likely promotes interaction with negatively charged TOM22 (isoelectric point ~4).^27^ The anionic character of TOM22 is instrumental for protein translocation across mitochondria, as translocating proteins contain positively charged pre-sequences. Previous studies identified a TOM22-binding peptide, KTGALLLQ,^28^ which is cationic and rich in aliphatic residues, like the sequences identified in our selection. Another TOM22-binding sequence identified in the same study, LCTKVPEL, contains 1 cationic and 1 anionic residue, and a balance of hydrophobic and hydrophilic amino acids, similar to cyclo[M-PELNRAI-K] identified in this work. This finding highlights the diversity of peptides targeting TOM22.

**Figure 5.**
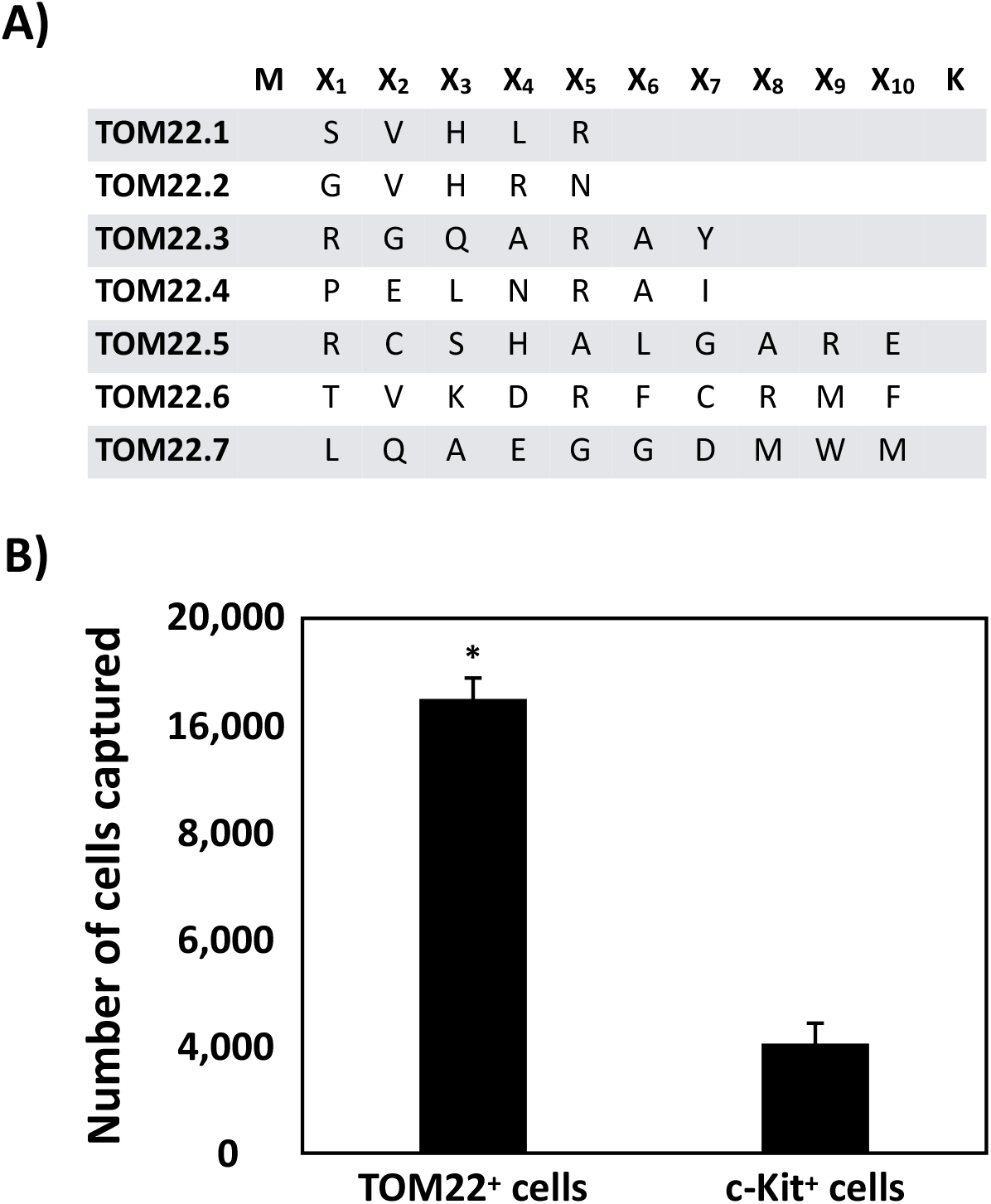
Identified TOM22-binding cyclic peptides and specificity characterization. **(A)** Amino acid sequences of peptides isolated from selections of a mRNA-display library of cyclic peptides against yeast-displayed TOM22. **(B)** Recovery of yeast cells displaying TOM22 or c-Kit by magnetic beads functionalized with cyclo[M-PELNRAI-K]. Yeast cells expressing TOM22 or c-Kit also displayed an engineered luciferase protein as a yeast surface fusion. Cell capture was quantified as a function of the luminescence signal produced by the recovered population using a generated luminescence standard curve. * represents p < 0.05 for a two tailed, paired t-test in comparison to the recovery of c-Kit displaying yeast cells.

We resolved to study the TOM22 binding of cyclo[M-PELNRAI-K]. The chemical diversity of this peptide, which contains positively (R) and negatively charged (E), polar (N), and aliphatic (A, I, L, and P) residues, is conducive to TOM22-binding by true affinity. Briefly, yeast cells co-expressing TOM22 and luciferase were incubated with magnetic beads functionalized with cyclo[M-PELNRAI-K] in the presence of non-displaying EBY100 yeast cells, which were included to reduce non-specific binding. Yeast cells bound to the peptide-functionalized beads were isolated using a magnet and subjected to a luminescence-based assay, wherein the luminescence signal is proportional to the number of TOM22-displaying cells captured by the magnetic beads. A study by Bacon *et al*. describes how this assay can be used to quantitatively rank ligands with different affinity for a target protein^29^; the correlation between the number of TOM22-displaying cells and luminescence signal is reported in a calibration curve detailed in **Figure S2**.

We also considered the binding of cells co-displaying a non-target protein (c-Kit) and a luciferase protein by the magnetic beads functionalized with peptide cyclo[M-PELNRAI-K]. We observed that TOM22^+^ yeast cells were captured by the peptide-functionalized beads at a statistically (~4.5-fold) higher level than TOM22^—^/c-KIT^+^ cells (**Figure 5B**), suggesting that cyclo[M-PELNRAI-K] is selective for TOM22. The luciferase-based assay indicates that the binding strength of the TOM22:cyclo[M-PELNRAI-K] complex is rather moderate, in the mid micromolar range. We therefore resolved not to evaluate this peptide in the context of targeted delivery to mitochondria. Nonetheless, the described luminescence-based assay was capable of detecting mild interactions owing to the multi-point binding mechanism (avidity) between cells and the peptide-functionalized beads, and the inherent sensitivity of the luciferase reporter.

Values of protein-binding affinity in the μM range, as those measured in this work, are typical for linear and cyclic peptides of 6-to-10 amino acids in length.^30^ It is however possible to enhance the affinity of peptide ligands selected combinatorially to reach a low-μM/high-nM range by adjusting the sequence and length of the protein-binding segment to increase the enthalpic component of the binding energy.^30^ Care should be taken in adjusting the protein-binding segment of cyclic peptide ligands to prevent excessive flexibility of the peptide backbone, which would unfavorably affect the entropic component of the binding energy. The library screening process can also be revisited to improve the binding affinity of isolated mutants by increasing the stringency of the selections. In this work, the selection conditions were maintained throughout the successive screening rounds, and only the elution conditions were adjusted; alternatively, competitive conditions can be adopted when incubating the library with the target cells by adding a mixture of soluble competitors, or by adjusting the composition, concentration, and pH of the selection buffer. Additional wash steps can also be performed to eliminate weak-affinity binders. Further, while held constant in this work, the number of protein-displaying cells could be reduced through the successive screening rounds to bias the isolation of higher affinity mutants. Finally, a higher number of selection rounds, beyond the four or five employed here, up to ten, can be performed to attain strong sequence homology among the identified peptides;^31^ nonetheless, this work has shown that specific binding peptides can be identified using fewer rounds of screening.

Collectively, our results show that the use of magnetic yeast displaying a target protein proves an ideal tool to increase the throughput of library screening, specifically for mRNA display libraries. It limits the need for recombinant soluble protein and provides an alternative screening format in the presence of a complex yeast surface. Improving the stringency of selection is nonetheless required to isolate high affinity binders from mRNA-display libraries due to the number of randomized amino acids included in the library backbone. Despite this limitation, the affinity of peptides identified in this study are comparable to other well-known WW binding peptides. Of interest also is that the lead candidate peptides performed better in their linear conformation as opposed to their cyclic conformation, which could be attributed to incomplete cyclization activity during the library synthesis. The experimental observation was also confirmed via molecular docking simulations. Moreover, the system was extended to a second protein of interest, TOM22, which gave an opportunity to showcase how ligands can be identified more efficiently for extracellular membrane proteins by selecting against magnetic yeast targets. Finally, a luminescence-based binding assay was used to rapidly confirm and quantify peptide:protein binding without target protein production. Development of this method and the successful identification of protein-binding peptides for two targets indicates that this platform, using yeast-displayed target proteins, could be used as to an alternative approach to screen mRNA display libraries. However, significant stringency is required to isolate high affinity binders.

## Associated Content

### Supporting Information

Supporting Information is available, which contains detailed experimental procedures, the DNA sequences used in this work, and supporting figures (PDF).

### Author Information

#### Corresponding Authors

**Stefano Menegatti** – Department of Chemical and Biomolecular Engineering, Biomanufacturing Training and Education (BTEC), North Carolina State University, Raleigh, 27695, North Carolina, United States.

**Balaji M Rao** – Department of Chemical and Biomolecular Engineering, Biomanufacturing Training and Education (BTEC), North Carolina State University, Raleigh, 27695, North Carolina, United States.

#### Authors

**John Bowen** – Department of Chemical and Biomolecular Engineering, Biomanufacturing Training and Education (BTEC), North Carolina State University, Raleigh, 27695, North Carolina, United States

**Kaitlyn Bacon** – Department of Chemical and Biomolecular Engineering, Biomanufacturing Training and Education (BTEC), North Carolina State University, Raleigh, 27695, North Carolina, United States

**Hannah Reese** – Department of Chemical and Biomolecular Engineering, Biomanufacturing Training and Education (BTEC), North Carolina State University, Raleigh, 27695, North Carolina, United States

#### Notes

The authors declare no competing financial interest.

## Acknowledgements

This work was funded by two grants from the National Science Foundation (CBET 1511227, CBET 1510845, and 1743404). KB and HR kindly acknowledge support from an NSF Graduate Research Fellowship. KB and JB kindly acknowledge support from a National Institute of Health Molecular Biotechnology Training Fellowship (NIH T32). We would also like to thank the UNC High-Throughput Peptide Synthesis and Array facility for peptide synthesis.

## Notes

### Competing Interest Statement

The authors have declared no competing interest.

