## Supplemental table and figures for "Use of target-displaying magnetized yeast in screening mRNA display peptide libraries to identify ligands"

#### Supplemental Methods

***Plasmids and yeast culture.*** The pCTCON and pCT302 vectors containing TRP or LEU selectable markers, respectively, were used in conjugation with *Saccharomyces cerevisiae* strain EBY100. Specifically, the Frozen-EZ yeast transformation Kit II (Zymo Research) was used to transform plasmid DNA into chemically competent EBY100. Trp-deficient SDCAA and SGCAA media was used, respectively, for culturing and inducing cells harboring the pCTCON plasmid.[1] Similarly, cells harboring pCT302 plasmids were grown and induced with Leu-deficient SDSCAA (-Leu) and SGSCAA (-Leu) media.[2] Both Trp-deficient and Leu-deficient media have similar compositions. However, the Leu-deficient media does not contain casamino acids, but rather uses synthetic dropout mix (1.62 g/L; US Biological Life Sciences) lacking leucine. When culturing, yeast cells were grown in SDCAA or SDSCAA media at 30°C with shaking at 250 RPM. To induce protein expression, yeast cells were transferred into SGCAA or SGSCAA medium at an $OD_{600}$ of 1 and incubated overnight at 20 °C with shaking at 250 rpm.

***Plasmid construction for display of proteins as yeast surface fusions and recombinant WW-YAP production.*** Plasmids were constructed that afford the expression of the proteins used for positive and negative selections of the mRNA display libraries as yeast surface fusions. These proteins were encoded as fusions to Aga2, a yeast cell mating protein. pCTCON-TOM22, affording the expression of TOM22, was constructed by amplifying gene block 1 with Pf1 and Pr1. Similarly, pCTCON-hFc, affording the expression of hFc (Fc portion of human IgG), was constructed by amplifying gene block 2 with primers Pf2 and Pr2. DNA amplified from gene blocks 1 and 2 were inserted into pCTCON between the NheI and BamHI cut sites. Finally, pCTCON-YAP-WW, affording the expression of the WW domains of YAP was constructed by amplifying gene block 3 with primers Pf3 and Pr3. DNA amplified from gene block 3 was inserted between the NheI and XhoI sites of pCTCON. The plasmid, pET22b(+), was utilized for recombinant WW-YAP protein expression. Gene block 3 was amplified with Pf4 and Pr4 and inserted between the NdeI and XhoI sites of pET22b(+), affording the expression of the soluble WW-YAP protein by creation of plasmid pET22b(+)-WW-YAP.

Double stranded gene fragments were purchased from Integrated DNA technologies (IDT). Primer oligonucleotides were bought from IDT or Eton Biosciences. Gene fragment and primer sequences can be found in **Tables S3-S4**. Phusion Polymerase (Thermo Fisher Scientific) was used for PCR reactions that took place in a 50 μL volume reaction following the manufacturer's protocols. Restriction digests of plasmid backbones and PCR products were executed at 37°C for 2 hours in a 50 μL volume using a 5-times excess of each restriction enzyme. Digested plasmid backbones were incubated with Antarctic phosphatase (New England Biolabs) for 1 hour at 37°C. Digested plasmids and PCR products were purified using a 9K series gel and PCR extraction kit (BioBasic). Overnight ligations using T4 DNA ligase (Promega) were performed with the digested plasmid backbones and PCR product inserts. Ligations were transformed into chemically competent Novablue *E.coli* cells. The cells were made chemically competent using Mix&Go! *E.coli* transformation buffers (Zymo Research). The GeneJET$^{TM}$ plasmid miniprep kit (Thermo Fisher Scientific) was used to harvest plasmids from overnight *E.coli* cultures.

***Construction of the cyclic peptide mRNA-display libraries.*** mRNA display libraries of randomized peptides were constructed using previously described protocols for guidance.[3,4] Oligonucleotides 1-3 **(Table S5)** encoding either a pentapeptide ($MX_1X_2X_3X_4X_5K$), heptapeptide ($MX_1X_2X_3X_4X_5X_6X_7K$), or decapeptide ($MX_1X_2X_3X_4X_5X_6X_7X_8X_9X_{10}K$) library were PCR-amplified using primers Pf5 and Pr5 **(Table S3)**. X represents any of the 20 amino acids. All oligonucleotides and primers were purchased from Integrated DNA Technologies. These primers add a 5' consensus sequence containing a T7 RNA polymerase promoter, a TMV translation enhancer, and a sequence coding the FLAG epitope tag. Also added is a 3' consensus sequence including a 6xHis tag followed by a sequence that affords conjugation of a puromycin linker. 25 PCR reactions were performed in a volume of 50 μL that contained 1 U of Phusion HF DNA polymerase (Thermo Fisher Scientific), 1X Phusion Buffer, 0.2 mM deoxynucleotide triphosphate (dNTPs) (Thermo Fisher Scientific), 0.1 μM of the forward and reverse primers, 1 M betaine, 3% dimethyl sulfoxide (DMSO), and 20 ng of the template DNA. The PCR was performed using the following conditions: Initial denaturation at 98°C for 2 minutes, followed by 30 cycles of denaturation at 98°C for 1 minute, annealing at 66°C for 1 minute, extension at 72°C for 15 seconds, and a final extension at 72°C for 10 minutes.

After, EDTA was added to the pooled PCR reactions at a final concentration of 5.5 mM. The amplified DNA was purified using phenol: chloroform: isoamyl alcohol precipitation followed by ethanol precipitation. Briefly, the DNA was extracted 2X with 1 volume of phenol: chloroform: isoamyl alcohol followed by an extraction with 1 volume of chloroform. After, 0.1 volumes of 3M potassium acetate, 2 volumes of ethanol, and 10 μL of linear acrylamide (Invitrogen) were added to the extracted DNA prior to overnight incubation at -20°C. The following day, the precipitated DNA was pelleted and washed 1X with 75% ethanol followed by a wash with 100% ethanol. After, the pellet was air dried prior to resuspension in 0.1% DEPC water.

The purified DNA was then used in an *in vitro* transcription reaction. The 300 μL reaction contained 12 μg of DNA, 5 mM ribonucleotide triphosphate (rNTPs) (Promega), 19 mM $MgCl_2$, 1X transcription buffer, 45 μL of T7 RNA polymerase (Thermo Fisher Scientific), and 0.1% DEPC water to reach the final volume. The reaction was incubated for 8 hours at 37°C. EDTA was added to a final concentration of 50 mM. After, the mRNA was purified by acidic phenol chloroform extraction (2X, 1 volume) followed by an extraction with chloroform (1X, 1 volume). The extracted mRNA was further purified using a Nap 5 column (GE Healthcare Life Sciences) by eluting in 0.1% DEPC water. Subsequently, the template DNA was digested in an 850 μL reaction for 4 hours at 37°C with 42.5 μL of RNase-free Turbo DNASE (1 U/μL, Thermo Fisher Scientific) and 1X DNase buffer. After, EDTA was added to a final concentration of 20 mM. The reaction mixture was purified using acidic phenol chloroform extraction followed by ethanol precipitation as previously described. This ethanol precipitation used lithium chloride instead of potassium acetate.

A puromycin linker ([psoralen-(ATAGCCGGTG)$_2$-OMe-dA$_{15}$-C9C9-Acc-puromycin]; Keck Oligo Synthesis Lab, Yale University) was conjugated to the purified mRNA. The conjugation reaction contained 200 μg of purified mRNA, 20 mM HEPES, 100 mM KCl, and the puromycin linker at 2.5 times the total molar concentration of mRNA. The total reaction volume was 250 μL. Conjugation took place in a thermocycler using the following conditions: 85°C for 8 minutes, then 60 cycles with a 1°C decrease in each cycle from 85°C to 25°C, followed by 25°C for 25 minutes, and a 4°C hold. Next, the puromycin linker was crosslinked to the mRNA using ultraviolet light (360 nm) for 20 minutes. The crosslinked mRNA was precipitated overnight using lithium acetate and ethanol. Prior to *in vitro* translation, the precipitated crosslinked mRNA was recovered via centrifugation and washed as previously described.

After, a 500 μL translation reaction was set up containing the crosslinked mRNA, 340 μL of rabbit reticulocyte lysate (Invitrogen), 50 μM methionine, 1X buffer without methionine, and nuclease free water to reach the final volume. The reaction mixture was incubated at 30°C for 1.5 hours prior to the addition of 1 M MgCl2 and 1 M KCl to final concentrations of 76 mM and 880 mM, respectively. After, this mixture was held at room temperature for 1 hour and then stored overnight at -20°C.

The puromycin linker contains a poly(dA) sequence allowing for purification of mRNA-puromycin and mRNA-puromycin-peptide fusions using oligo dT beads. The translation mix was added to 1 mL of magnetic oligo-dT beads (New England BioLabs) that had been washed prior 3X with 0.1% DEPC water and 3X with 10 mM Tris pH 7.5, 1 mM EDTA, and 0.05% SDS. The translation mix and washed beads were incubated in 9 mL of fresh binding buffer (0.5 M NaCl, 10 mM Tris pH 7.5, 1 mM EDTA, 0.05% SDS, and 1 mM DTT) for 2 hours at 4°C. Next, the beads were washed 3x10 minute with wash buffer (0.2 M NaCl, 10 mM Tris pH 7.5, 1 mM EDTA, 0.05% SDS, 1 mM DTT) followed by washes 3x10 minutes with crosslinking buffer (0.2 M NaCl, 1 mM EDTA, 0.05% SDS, 1 mM DTT). Subsequently, cyclization was performed by resuspending the beads in 800 μL of crosslinking buffer and adding 50 μL of a 3 mg/mL solution of a disuccinimidyl glutarate crosslinker (Thermo Fisher Scientific) dissolved in dimethylformamide. The cyclization reaction took place for two hours at 4°C. After, the beads were washed 3x10 minute with crosslinking buffer. The cyclization reaction was repeated a second time. Lastly, the beads were washed 3x10 minute with wash buffer and then eluted in 600 μL of 0.1% DEPC water with 1 mM dTT overnight at 4°C.

The cyclized oligo(dT) purified product underwent a reverse transcription reaction the following day. Initially, 25 μL of 100 μM of a reverse transcription primer (5'- TTT TTT TTT TNN CCA GAT CCA GAC ATT CCC AT-3') was incubated with the oligo(dT) purified product for 15 minutes at room temperature. Next, 200 μL of 5X first strand buffer, 50 μL of 10 mM dNTPs, 100 μL of 0.1 M DTT, and 20 μL of 0.1% DEPC water was added to the reaction mixture. The reaction mixture was then incubated at 42°C for 2 minutes prior to the addition of 5 μL of Superscript™ II reverse transcriptase (Invitrogen). Subsequently, the reaction was held for 50 minutes at 42°C. After, EDTA was added to the reaction at a final concentration of 6 mM. The reaction mixture was then passed through a Nap-10 column (GE Healthcare Life Sciences) using 20 mM Tris pH 7.5, 150 mM NaCl, 0.05% Tween 20 as the equilibration and elution buffers.

Magnetic Ni-NTA agarose beads (Thermo Fisher Scientific, Pierce) were used to isolate the mRNA-cDNA-peptide fusions by taking advantage of the 6xHis tag on the translated peptide. 1 mL of beads was washed with binding buffer (50 mM Sodium Phosphate pH 8, 300 mM NaCl).

After, the Nap-10 column eluate was incubated with the beads at room temperature for 1 hour. The beads were washed 3X with binding buffer and eluted with 500 μL of 50 mM sodium phosphate pH 8, 300 mM NaCl, 300 mM imidazole. Subsequently, the eluted mRNA-cDNA-peptide fusions were desalted using a Nap-5 column by eluting with 0.1% DEPC water. Library diversity was estimated using a $A_{260}$ measurement.

***Magnetization of yeast cells.*** $5x10^7$ yeast cells expressing each selection protein were incubated in 2 mL of their respective magnetization buffer with 0.4 mg iron oxide (4 mg/mL in water) for 30 minutes at room temperature. Yeast cells displaying TOM22 were magnetized in 50 mM sodium acetate pH 5, 50 mM NaCl buffer. Yeast cells displaying WW-YAP and hFc were magnetized in 50 mM Tris-HCl pH 7.4, 300 mM NaCl buffer. Different buffers were used as it appeared iron oxide binding was dependent on the isoelectric point of the yeast displayed target protein. Any cells bound to the iron oxide were isolated from the unbound cells using a magnet. The $OD_{600}$ of the solution prior to the addition of the iron oxide ($OD_I$) and after the removal of the yeast bound to the iron oxide ($OD_f$) was measured. Yeast recovery was calculated as $(OD_I - OD_f)/OD_f$. ~ $2x10^7$ cells were magnetized and used in each screen.

After the initial magnetization, the yeast-iron oxide conjugates were washed three times to remove any unbound or weakly bound cells using the appropriate magnetization buffer. Subsequently, the yeast-iron oxide conjugates were incubated with PBS pH 7.4, 1% BSA, 0.1% salmon sperm DNA (1% PBSASD) for 1 hour at room temperature to block any unbound iron oxide. After blocking, the yeast-iron oxide conjugates were washed two more times with 1% PBSASD to remove any yeast cells that may have dissociated from the iron oxide. After these steps, the magnetic yeast were used as targets in the selection of mRNA display libraries.

***mRNA-display screening to identify affinity peptides using magnetic yeast targets.*** A cyclic peptide library with five randomized amino acid positions was used to screen for peptide binders to WW-YAP or TOM22. Four or five rounds of screening were performed when identifying ligands to WW-YAP and TOM22, respectively. During each round, a negative selection was initially performed against yeast cells expressing hFc. The mRNA-cDNA-peptide fusions were incubated with magnetic yeast cells displaying hFc in 1 mL of PBS pH 7.4, 0.1% BSA, 0.1% salmon sperm DNA for 1 hour at room temperature. Subsequently, the incubation mixture was placed on a magnet and the supernatant containing unbound mRNA-cDNA-peptide fusions was removed and incubated with magnetic yeast cells displaying either WW-YAP or TOM22. After an additional 1-hour incubation, the selection mixture was placed on a magnet and the supernatant was removed to isolate any mRNA-cDNA-peptide fusions positively bound to the target displaying cells.

The selection stringency was increased for each round of screening. For round one, peptide-yeast-iron oxide conjugates were not washed after separation using a magnet. The bound peptide fusions were eluted twice in 200 μL of 0.15 M potassium hydroxide for 1 hour. The eluted fractions were combined, and the cDNA associated with the eluted fusions was used as a template to generate the library for the next screening round. For round two, the elution steps were similar, but the bound peptides fusions were washed 3 times with gentle pipetting using 0.1% BSA PBS (0.1% PBSA) prior to elution. For round three, the same wash steps were carried out, but the peptides fusions were first incubated in 200 μL of a weak, low pH buffer (50 mM sodium acetate, 50 mM NaCl pH 5) for 1 hour at room temperature, followed 2X elution in 0.15 M potassium hydroxide for 1 hour at room temperature. The library was remade for the next screening round with cDNA obtained after elution using potassium hydroxide to bias the isolation of higher affinity binders. For rounds four and five, the same washing and elution steps were carried out as in round three, but an additional hour of elution in the weak, low pH 5 buffer was performed. All eluates were neutralized using 5N HCl.

***Library reconstruction for subsequent screening rounds.*** For each elution condition, the mRNA-cDNA-peptide fusions eluted in each incubation were pooled together. For rounds three through five, only the fusions eluted using the harsher potassium hydroxide buffer were pooled and precipitated for DNA amplification. cDNA was precipitated using ethanol precipitation and linear acrylamide as previously described. The precipitated cDNA was used to generate a new mRNA display library for the subsequent screening round. The precipitated cDNA was amplified using primers Pf5 and Pr5. A 50 μL PCR reaction was performed containing 1 U Phusion HF DNA polymerase, 1X HF Phusion buffer, 0.2 mM dNTPS, 0.2 μM of each primer, 1 M betaine, 3% DMSO, and the precipitated cDNA template. The PCR was performed using the following conditions: Initial denaturation at 98°C for 2 minutes, followed by 30 cycles of denaturation at 98°C for 1 minute, annealing at 66°C for 1 minute, extension at 72°C for 15 seconds, and a final extension at 72°C for 10 minutes. The PCR product was ethanol precipitated using linear acrylamide as previously described and used as the template for additional PCRs. A visible product band was not observed on a gel until the second round of PCR. Additional PCR reactions were performed to obtain enough DNA for library generation. Similar steps as described were carried out using this amplified DNA to generate mRNA-cDNA-peptide fusions for the next round of screening.

After the final round of screening, the fusions eluted using potassium hydroxide were precipitated and amplified using similar conditions. The amplified DNA was inserted into the pJet2.1 vector via blunt-end ligation using the CloneJET kit (Thermo Fisher Scientific). The ligation mixture was transformed into chemically competent Novablue *E. coli* cells. DNA from 10 individual colonies was extracted using the GeneJET™ plasmid miniprep kit and sent for sequencing. Some isolated clones contained premature stop codons and were not considered.

***Synthesis of WW domain-binding linear and cyclic peptides.*** The linear peptides MAFRLCK, MLDFVNHRSRGK, MDGNLSGIMPVK, and MYRGDPETCVDK were synthesized on 0.6 mL of Toyopearl AF-Amino-650 M (corresponding to 0.1 g of dry resin) (Tosoh) following conventional Fmoc/tBu peptide synthesis.[5] All amino acid conjugations were performed using 2 equivalents (compared to the Toyopearl amine density of 0.2 meq/g) of Fmoc-protected amino acids (ChemImpex), Hexafluorophosphate Azabenzotriazole Tetramethyl Uronium (HATU, 2 eq.), and diisopropylethylamine (DIPEA, 2 eq.) in anhydrous N,N'-dimethylformamide (DMF); 2 conjugations per amino acid were performed at 75°C for 15 minutes under sonication using a Syro I automated peptide synthesizer (Biotage). Completion of the amino acid conjugation was qualitatively confirmed by Kaiser test (Millipore Sigma). Fmoc deprotection was performed by incubating the resin with 2 mL of 20%v/v piperidine (Millipore Sigma) in DMF. Following chain elongation, the peptides were deprotected by acidolysis using a 90:5:3:2 trifluoroacetic acid:thioanisole:ethane dithiol:anisole (Millipore Sigma) cocktail for 2 hours at room temperature under mild agitation.

The cyclic peptides cyclo[M-AFRLC-K], cyclo[M-LDFVNHRSRG-K], cyclo[M-DGNLSGIMPV-K], and cyclo[M-YRGDPETCVD-K] were synthesized on Toyopearl AF-Amino-650 M as follows. First, Fmoc-Lys(Mtt)-OH (2 eq.) was conjugated on Toyopearl resin using HATU (2 eq.) and DIPEA (2 eq.) in anhydrous DMF. The linear sequences MAFRLCK, MLDFVNHRSRGK, MDGNLSGIMPVK, and MYRGDPETCVDK were then synthesized as described above. Following Fmoc deprotection of methionine, the N-terminal amine was succinylated by reaction with succinic anhydride (6 eq.) and dimethylaminopyridine (DMAP, 3 eq.) in anhydrous DMF for 24 hours at room temperature under mild agitation. The Mtt protecting group was removed from the C-terminal lysine by incubation with a cocktail of 96:6:2 dichloromethane:trifluoroacetic acid:triisopropyl silane for 10 minutes at room temperature under mild agitation. The carboxyl group of the succinyl-peptide was activated with HATU (2.98 eq.) and DIPEA (6 eq.) in anhydrous DMF and incubated for 15 minutes at 75$^{o}$C to enable peptide cyclization by forming an amide bond with the ε-amine of a C-terminal lysine. The carboxyl activation and cyclization reactions were repeated in order to ensure a high yield of cyclized peptide. Following peptide cyclization, the peptides were deprotected by acidolysis as described above. All resins were sequentially washed with dichloromethane (DCM), DMF, and DCM, then dried under air flow, and finally stored at -20$^{o}$ following deprotection.

***Recombinant YAP-WW protein expression and purification.*** Plasmid pET22b(+)-WW-YAP was transformed into chemically competent Rosetta *E. coli* cells. A 5 mL culture was grown in LB media (10 g/L tryptone, 5 g/L yeast extract, 10 g/L NaCl) overnight at 37$^{o}$ C. Expression was carried out in 2XYT media (16 g/L tryptone, 10 g/L yeast extract, and 5 g/L NaCl). The 5 mL overnight culture was to inoculate a 1 L culture of 2XYT. The cells were induced at a concentration of 0.5 mM IPTG when the OD reached between 0.8 and 1.0. Induction occurred for 20 hours at 20$^{o}$ C.

The YAP-WW protein was purified by metal affinity chromatography as follows. Briefly, cells were pelleted via centrifugation for 12 minutes at 3,500xg and resuspended in 35 mL of resuspension buffer (20 mM HEPES, 150 mM NaCl pH 7.8, 0.2 mM phenylmethylsulfonyl). The cell slurry was sonicated, and the unlysed cells and cell debris was pelleted for 22 minutes at 15,000xg. The supernatant was then syringe filtered (Becton Dickinson) using a 0.45 µm PVDF (Genesee Scientific) filter. Filtered cell lysate was loaded onto a 5 mL Bio-Rad Nuvia IMAC column at 2 mL/min, washed with 50 mL of Buffer C-IMAC (20 mM HEPES, 800 mM NaCl pH 7.8) at 5 mL/min, flushed with 50 mL of Buffer A-IMAC (20 mM HEPES, 137 mM NaCl pH 7.8) at 5 mL/min, and eluted with a 50 mL linear gradient of 0-100% Buffer B-IMAC (20 mM HEPES, 137 mM NaCl, 500 mM Imidazole pH 7.8) at 5 mL/min. Chromatographic fractions were analyzed via SDS-PAGE, and fractions containing the protein of interested were pooled. Pure fractions were subsequently dialyzed into 20 mM HEPES, 150 mM NaCl pH 7.8 (HEP) using SnakeSkin™ Dialysis Tubing (MWCO 3.5 kDa) (Thermo Fisher Scientific). The protein was held at -80$^{o}$C in HEP plus 10% glycerol for long-term storage.

***Binding and elution of WW-YAP using peptide-Toyopearl resins.*** 30 mg of functionalized Toyopearl Amino resin was swelled overnight in 20% methanol at 4$^{o}$C. Each swelled resin was washed 3X with PBS the following day prior to incubation with WW-YAP. Frozen aliquots of recombinant WW-YAP protein were thawed at room temperature prior to dilution in PBS. 0.4 mg of WW-YAP (1 mg/mL) was incubated with 15 mg of resin for 1 hour at room temperature. After, the supernatant was isolated. The amount of protein present in the supernatant was quantified via an $A_{280}$ absorbance measurement. The amount of protein bound by the resin was calculated via mass balance, and the amount of bound protein was recorded as a percentage of the initial total protein.

Subsequently, a bind and elute study was executed to evaluate the top two binding candidates. 100 mg of Toyopearl amino resin functionalized with each peptide candidate was swelled overnight in 20% methanol at 4°C. Prior to incubation with WW-YAP, each swelled resin was washed 3X with HEP. 0.2 mg of WW-YAP (1 mg/mL) was incubated with 5 mg of resin for 1 hour at room temperature. After, the supernatant was removed, and the resin was with washed 1X with HEP. The resin was incubated with 200 µL of 100 mM glycine pH 2.0 for 5 minutes at room temperature to elute the bound protein. The unbound, wash, and eluted fractions were analyzed via $A_{280}$ absorbance measurements to determine the protein content.

***Determination of binding capacity and affinity of WW-YAP using peptide-Toyopearl resins via isotherm adsorption study.*** WW-YAP binding isotherms were constructed for the top two lead peptides as follows. A frozen stock of WW-YAP protein was diluted with HEP as appropriate. 30 mg of functionalized Toyopearl Amino resin was swelled overnight in 20% methanol at 4°C. The resin was washed 3X with HEP prior to incubation with WW-YAP. WW-YAP at varying concentrations (20 µg/mL to 12.5 mg/mL) in a volume of 400 µL was incubated with 1 mg of functionalized resin for 1 hour at room temperature. The supernatant was removed and an $A_{280}$ absorbance measurement of the supernatant was performed for protein quantification. The amount of protein bound to the resin was calculated via mass balance. The $K_D$ describing the binding interaction between the peptide-functionalized Toyopearl resin and soluble WW-YAP protein was determined using the relationship:

$$q = \frac{q_{max}[C]}{K_D + [C]} \ \textbf{(Eq. 1)}$$

where $q$ is the amount of protein bound to the resin (mg protein/mL resin), $q_{max}$ is the maximum protein binding capacity (mg protein/mL resin), *[C]* is the unbound concentration of protein at equilibrium (mg protein/mL solution), and $K_D$ is the binding affinity constant (mg protein/mL solution). A global non-linear least squares regression was used to fit the data to **Eq. 1**.[1] A single $K_D$ and $q_{max}$ value were fit to the binding data across 3 repeats where the soluble protein concentration ranged from 0-2 mg/mL.

A combination of a Langmuir isotherm and a Freundlich isotherm was also used to fit the entire soluble protein concentration range from 0-10 mg/mL using the relationship

$$q = \frac{q_{max}[C]}{K_D + [C]} + K[C]^{\frac{1}{n}} \ \textbf{(Eq. 2)}$$

where $K$ is a dimensionless distribution coefficient and $n$ is a dimensionless correction factor. A global non-linear least squares regression was used to fit the data to **Eq. 2**. A single $K_D$, $q_{max}$, $K$, and $n$ value were fit to the binding data across 3 repeats where the soluble protein concentration ranged from 0-12.5 mg/mL.

***In silico modeling of WW:peptide interactions.*** The crystal structures of the WW domain of human YAP (PDB: 2LTW)[6] was subjected to standard protein preparation using Schrödinger's ProteinPrep Wizard[7,8] to search for and correct missing atoms and/or entire side chains (with the PRIME software), remove extra salts and non-binding ligands, add explicit hydrogens, assign tautomeric states with EPIK, optimize hydrogen bonding networks, and minimize the protein's energy with the OPLS3e force field.[9]

Peptides MAFRLCK-GSG, MLDFVNHRSRGK-GSG, cyclo[M-AFRLC-K]-GSG, and cyclo[M-LDFVNHRSRG-K]-GSG were initially designed using the molecular editor Avogadro[10,11] and were equilibrated by molecular dynamics in the GROMACS[12–14] simulation package using the OPLS all-atom force field[15–17] and periodic boundary conditions[18–20]. The peptide was individually placed in a simulation box with periodic boundary conditions containing 800 water molecules (TIP3P water model).[21] The solvated system was initially minimized by running 10,000 steps of steepest gradient descent, heated to 300 K in an NVT ensemble for 250 ps with 1 fs time steps, and equilibrated to 1 atm by running a 500-ps NPT simulation with 2 fs time steps. The production run for every peptide was performed in the NPT ensemble, at constant T = 300 K using the Nosé-Hoover thermostat[22–24] and constant P = 1 atm using the Parrinello-Rahman barostat[25,26]. The coordinates of atoms were saved every 2ps. The leap-frog algorithm was used to integrate the equations of motion, with integration steps of 2 fs, and all of the covalent bonds were constrained by means of the LINCS algorithm[27]. The short-range electrostatic and Lennard-Jones interactions were calculated within a cutoff of 1.0 nm and 1.4 nm, respectively, whereas the particle-mesh Ewald method was utilized to treat the long-range electrostatic interactions.[28–30] The non-bonded interaction pair-list was updated every 5 fs, using a cutoff of 1.4 nm.

The peptides were then docked *in silico* against putative binding sites on the WW domains of human YAP using the docking software HADDOCK (High Ambiguity Driven Protein-Protein Docking, v.2.2).[31–33] Default HADDOCK parameters (*e.g.*, temperatures for heating/cooling step and number of molecular dynamics sets per stage) were used. All the residues on each binding site (solvent accessibility of 50% or greater) were defined as "active", whereas the residues surrounding the binding sites were defined as "passive". All variable amino acid positions on the peptide ligands were also denoted as 'active', while the GSG tripeptide spacer was defined as not being involved in the interaction to account for the directionality of binding. Docking proceeded through a 3-stage protocol: (1) rigid, (2) semi-flexible, and (3) water refined fully flexible docking. A total of 1000, 200, and 200 structures were calculated at each stage, respectively. Final structures were grouped using a minimum cluster-size of 20 (10% of the total water refined calculated structures) with a Cα RMSD < 7.5 Å using ProFit (Martin, A.C.R, http://www.bioinf.org.uk/software/profit/). Once the clusters were identified for each WW:peptide complex pair, FireDock[34,35] and XScore[36,37] were used to score the complexes; FireDock is an efficient method re-scoring of protein-protein docking solutions, while Xscore computes the dissociation of a protein-ligand complex using an empirical equation that considers energetic factors in the protein-ligand binding process. The selected binding poses were then refined via 100-ns atomistic molecular dynamics (MD) simulations using the Gromacs simulation package. The WW:peptide complexes were embedded in a cubic periodic box of 9.7 nm side lengths and solvated with 30,000 TIP3P water molecules. The MD simulations were performed at 300 K and 1 atm using the Amber99SB force field. The MM/GBSA method was used for post-processing of the WW:peptide complexes derived from MD simulations[38,39]; if poses had conflicting docking score or MM-GB/SA ranks, then the pose was discarded.

***Specificity analysis of a TOM22 binding peptide using a luminescence binding assay.*** The specificity of cyclo[M-PELNRAI-K] for TOM22 was evaluated using a luminescence based binding assay that takes advantage of a luciferase reporter to quantify recovery.[40] We used plasmids that afford the co-expression of a target protein and NanoLuc, an engineered luciferase. Previously described plasmid pCT302-T2A-NanoLuc was digested between its NheI and BamHI sites. Gene block 1 encoding the DNA for TOM22 was amplified using primers Pf1 and Pr1 and inserted into the digested plasmid backbone to create pCT302-TOM22-T2A-NanoLuc. Similarly, Gene block 4 encoding the DNA for the extracellular domain of c-Kit was amplified using primers Pf6 and Pr6 and inserted into the digested plasmid backbone to construct pCT302-cKit-T2A-NanoLuc.

Peptide cyclo[M-PELNRAI-K] was synthesized in its cyclic form by the UNC High-Throughput Peptide Synthesis and Array facility. The synthesized peptide included a C-terminal biotin and was cleaved from resin. The peptide was purified by RP18 HPLC (purity 95%) and its mass was confirmed by MALDI mass spec.

cyclo[M-PELNRAI-K] was immobilized onto the surface of magnetic streptavidin beads. Briefly, 25 μL of Magnetic biotin binder Dyanbeads™ (Invitrogen) were washed 2X with 0.1% PBSA. After, the beads were incubated with 11 μM of the TOM22 binding peptide overnight in a total volume of 100 μL using 0.1% PBSA at 4°C. The next morning the beads were washed 3X with 0.1% PBSA followed by an hour incubation in PBS pH 7.4, 1% BSA for blocking. The beads were then washed 2X with 0.1% PBSA, 0.05% tween-20 (0.1% PBSAT). Subsequently, the functionalized beads were incubated with either $1x10^7$ cells co-expressing TOM22 and NanoLuc or c-Kit and NanoLuc in a total volume of 2 mL using 0.1% PBSAT for 1 hour at room temperature. Also included were $1x10^9$ EBY100 cells to reduce non-specific binding. When the incubation was complete, the beads were washed 3X with 0.1% PBSAT and then resuspend in 100 μL of PBS.

The binding of the cells to the functionalized magnetic beads was detected using the Nano-Glo Luciferase Assay system (Promega). 100 μL of the reconstituted reagent was incubated with the magnetic beads. The reaction proceeded for 3 minutes. After, the tube was placed on a magnet and 100 μL of the supernatant was plated onto a 96 white-well plate with a clear bottom (Corning) in duplicate. A Tecan Infinite 200 Plate Reader was used to read the luminescence signal with the following settings: integration time of 1000 ms, settle time of 0 ms, and no attenuation.

Calibration curves were made for the TOM22 and c-Kit reporter cells to develop a relationship between luminescence signal and number of reporter cells incubated. A known number of reporter cells ranging from $1x10^5$ - $1x10^7$ were resuspended in 100 μL of PBS. Luminescence assays were carried out as previously described for each concentration of cells to determine the points in the calibration curve. The luminescence of an equal volume of PBS and the Nano-Glo Luciferase Assay Reagent was also considered as a blank. Background subtracted luminescence signals were plotted against the number of reporter cells incubated to generate a standard curve. A linear regression was also fit to the data points. For each pull down assay, the fitted linear regression was used to estimate the number of reporter cells captured by the peptide functionalized beads using the luminescence signal produced by the captured cells.

### Supplemental Figures

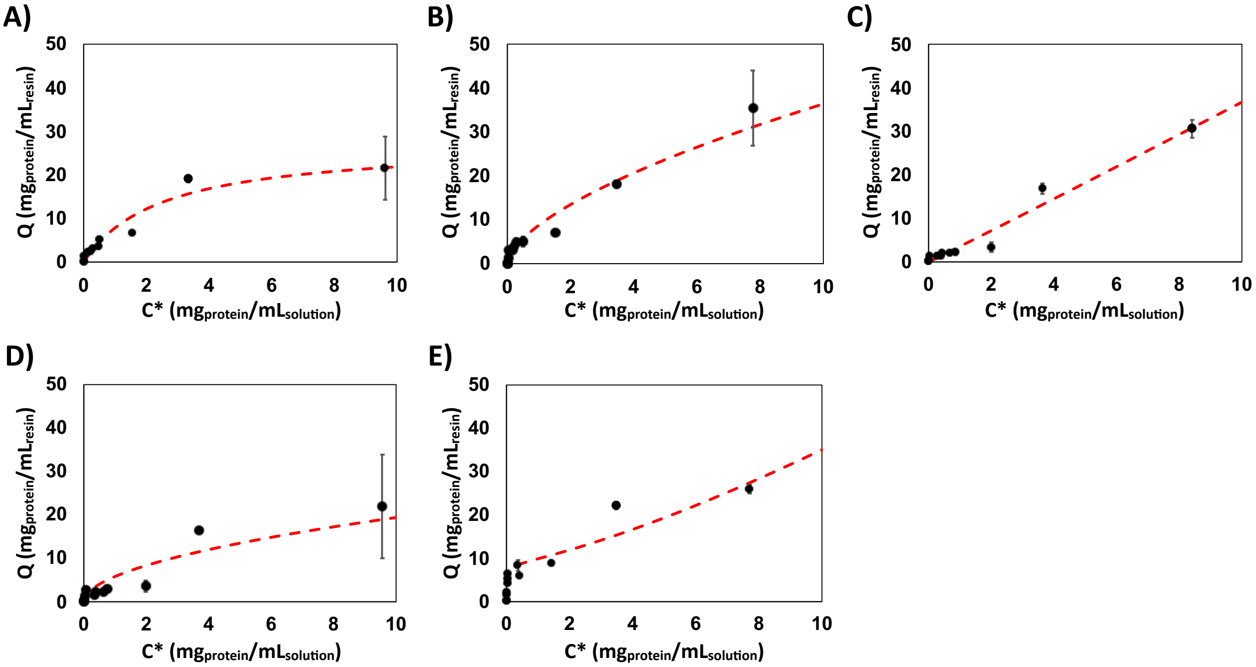


**Figure S1.** WW-YAP equilibrium binding isotherms for **(A)** cyclo[M-AFRLC-K]-Toyopearl, **(B)** MAFRCLK-Toyopearl, **(C)** cyclo[M-LDFVNHRSRG-K]-Toyopearl, **(D)** MLDFVNHRSRGK-Toyopearl, and **(E)** RYSPPPPYSSHS-Toyopearl. The binding isotherms were generated by fitting the values of protein bound to the resin (*Q*) *vs.* the concentration of protein in solution at the equilibrium (*C**) against a combined Langmuir and Freundlich isotherm model. The combined model was fit across WW-YAP concentrations ranging from 20 µg/mL to 10 mg/mL.

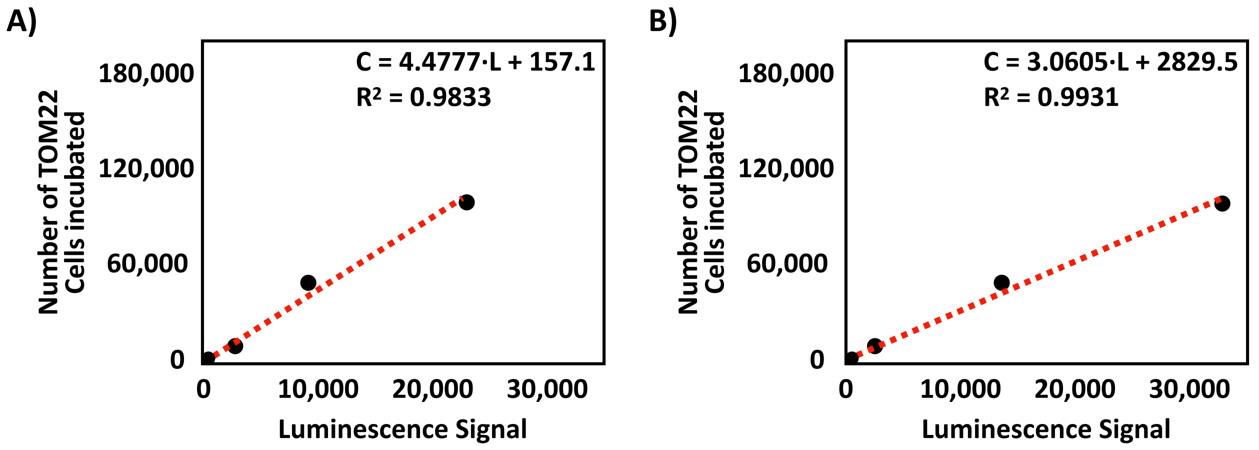


**Figure S2.** Example of calibration curves used to correlate luminescence signal to the number of TOM22 (**A**) or c-Kit (**B**) cells incubated. These cells also co-expressed a luciferase protein. A new calibration curve specific to each cell type was generated for each repeat. A linear regression was fit to the data to relate the number of cells present (C) to the produced luminescence signal (L). This relationship was used to estimate the number of TOM22 or c-Kit cells captured by the peptide functionalized magnetic beads based on the luminescence signal produced by the isolated cells.

**Table S1.** Values of affinity ($K_D$), maximum binding capacity ($Q_{max}$), distribution coefficient ($K$), and correction factor ($n$) obtained from the combined Langmuir-Freundlich fits in **Figure S1**.

| Sequence | $K_D$ (mg/mL) | $Q_{max}$ (mg/mL$_{resin}$) | K | n | $K_D$ (µM) |
|---|---|---|---|---|---|
| **cyclo[M-AFRLC-K]** | 2.4780 | 27.26 | 0.00 | 2.42 | 184.9 |
| **M-AFRLC-K** | 0.0697 | 0.00 | 8.77 | 1.62 | 5.2 |
| **cyclo[M-LDFVNHRSRG-K]** | 0.8730 | 0.00 | 3.55 | 0.99 | 65.2 |
| **M-LDFVNHRSRG-K** | 0.0472 | 0.00 | 5.81 | 1.91 | 2.7 |
| **PTCH peptide** | 0.0381 | 8.99 | 1.24 | 0.76 | 2.8 |

**Table S2.** Values of WW-YAP recovery from bind-and-elute studies using cyclo[M-AFRLC-K]-Toyopearl and cyclo[M-LDFVNHRSRG-K]-Toyopearl resins. Each resin was incubated with ~1.2 mg/mL WW-YAP.

| Sequence | WW-YAP Recovery (%) | 68% Confidence Interval | Mass Balance Closure (%) | 68% Confidence Interval |
|---|---|---|---|---|
| **cyclo[M-AFRLC-K]-Toyopearl** | 87 | 82 - 93 | 94 | 91 - 96 |
| **cyclo[M-LDFVNHRSRG-K]-Toyopearl** | 78 | 75 - 82 | 93 | 92 - 95 |

**Table S3**. List of oligonucleotide primers.

| | Sequence |
|---|---|
| Pf1 | GTTCTCGCTAGCGCTGCCGCCGTCGCTG |
| Pr1 | GCACTTGGATCCTGCCCTGGAAAACCTGTACATTT |
| Pf2 | GTTGACGCTAGCCCCAAATCTTGTGACAAAACTCACA |
| Pr2 | GCACTTGGATCCTTTACCCGGAGACAGGGAGA |
| Pf3 | AAAAAAGCTAGCTTCGAAATTCCCGACGAT |
| Pr3 | AAAAAACTCGAGCTATTACAAGTCCTCTTCAGAAATAAGCTTTTGTTCCTGATTCATCGC |
| Pf4 | AAAAAACATATGTTCGAAATTCCCGACGATGTG |
| Pr4 | AAAAAACTCGAGCTGATTCATCGCAAAGCGTG |
| Pf5 | GCAAATTTCTAATACGACTCACTATAGGGACAATTACTATTTACAATTAC |
| Pr5 | ATAGCCGGTGCCAGATCCAGACATTCCCATATGGTGATGGT |
| Pf6 | GTTGACGCTAGCCAACCATCTGTGAGTCCAGGG |
| Pr6 | GCACTTGGATCCAGGAGTGAACAGGGTGTGGG |

**Table S4.** List of gene block fragments.

| | |
|---|---|
| Gene Block 1 | GCTGCCGCCGTCGCTGCTGCCGGTGCAGGGGAACCCCAGTCCCCGGACGAATTGCTCCCG<br>AAAGGCGACGCGGAGAAGCCTGAGGAGGAGCTGGAGGAGGACGACGATGAGGAGCTA<br>GATGAGACCCTGTCGGAGAGACTATGGGGCCTGACGGAGATGTTTCCGGAGAGGGTCCG<br>GTCCGCGGCCGGAGCCACTTTTGATCTTTCCCTCTTTGTGGCTCAGAAAATGTACAGGTTTT<br>CCAGGGCA |
| Gene Block 2 | CCCAAATCTTGTGACAAAACTCACACATGCCCACCGTGCCCAGCACCTGAACTCCTGGGGG<br>GACCGTCAGTCTTCCTCTTCCCCCCAAAACCCAAGGACACCCTCATGATCTCCCGGACCCCT<br>GAGGTCACATGCGTGGTGGTGGACGTGAGCCACGAAGACCCTGAGGTCAAGTTCAACTG<br>GTACGTGGACGGCGTGGAGGTGCATAATGCCAAGACAAAGCCGCGGGAGGAGCAGTACA<br>ACAGCACGTACCGTGTGGTCAGCGTCCTCACCGTCCTGCACCAGGACTGGCTGAATGGCA<br>AGGAGTACAAGTGCAAGGTCTCCAACAAAGCCCTCCCAGCCCCCATCGAGAAAACCATCTC<br>CAAAGCCAAAGGGCAGCCCCGAGAACCACAGGTGTACACCCTGCCCCCATCCCGGGATGA<br>GCTGACCAAGAACCAGGTCAGCCTGACCTGCCTGGTCAAAGGCTTCTATCCCAGCGACATC<br>GCCGTGGAGTGGGAGAGCAATGGGCAGCCGGAGAACAACTACAAGACCACGCCTCCCGT<br>GCTGGACTCCGACGGCTCCTTCTTCCTCTACAGCAAGCTCACCGTGGACAAGAGCAGGTG<br>GCAGCAGGGGAACGTCTTCTCATGCTCCGTGATGCATGAGGCTCTGCACAACCACTACACG<br>CAGAAGAGCCTCTCCCTGTCTCCGGGTAAA |
| Gene Block 3 | ATGGATCCTGGGCAGCAGCCGCCACCCCAACCAGCGCCTCAAGGTCAGGGGCAACCGCCA<br>TCACAGCCCCCTCAGGGCCAAGGACCTCCGTCAGGCCCTGGTCAGCCAGCGCCCGCAGCG<br>ACACAGGCCGCACCCCAGGCTCCGCCCGCAGGTCACCAGATAGTGCACGTTCGTGGAGAC<br>AGTGAAACCGACCTGGAAGCCCTGTTCAATGCGGTTATGAACCCTAAGACCGCTAATGTAC<br>CACAAACGGTGCCGATGAGACTTAGAAAACTTCCAGATAGCTTTTTCAAACCACCCGAGCC<br>CAAGTCTCATAGTCGTCAAGCTTCAACAGACGCAGGAACGGCCGGGGCGTTGACTCCTCA<br>GCACGTGCGGGCTCACTCGTCACCTGCCAGTTTACAGTTGGGTGCTGTCTCACCCGGTACG<br>CTGACCCCTACTGGCGTAGTGTCCGGTCCGGCCGCAACACCTACGGCCCAACACCTTCGTC<br>AATCATCGTTCGAAATTCCCGACGATGTGCCGCTGCCAGCCGGTTGGGAGATGGCCAAAA<br>CTTCCTCAGGCCAGCGTTACTTCTTAAACCACATAGATCAGACAACTACCTGGCAGGATCC<br>GCGGAAAGCGATGTTAAGTCAAATGAATGTGACGGCACCTACATCCCCGCCAGTCCAGCA<br>GAATATGATGAACTCTGCTAGTGGTCCTCTTCCTGATGGGTGGGAACAGGCGATGACGCA<br>AGACGGCGAAATTTATTACATAAACCACAAGAACAAAACAACCTCGTGGCTTGATCCACG<br>GCTGGACCCACGCTTTGCGATGAATCAGCGCATCTCTCAAAGTGCCCCAGTTAAACAACCA<br>CCGCCCTTAGCTCCGCAGTCGCCACAGGGTGGGGTGATGGGCGGCAGCAACTCGAATCAG<br>CAGCAGCAAATGCGCTTACAGCAACTTCAGATGGAAAAGGAACGCTTACGCTTGAAACAA<br>CAAGAGCTGTTGCGGCAAGAATTAGCATTAAGATCACAGCTGCCTACTTTAGAACAGGAC<br>GGGGGTACGCAAAATCCCGTATCCTCGCCGGGAATGAGTCAGGAGCTTCGTACTATGACA<br>ACGAACTCCTCTGATCCTTTCCTTAACTCCGGGACCTATCACTCGCGCGACGAAAGTACTGA<br>TTCCGGATTGTCCATGTCCTCCTATTCCGTCCCCCGGACCCCCGACGATTTCCTTAACTCGGT<br>TGACGAAATGGACACTGGGGATACTATAAACCAATCGACCCTGCCTAGCCAGCAAAACCG<br>TTTCCCGGATTATTTAGAAGCTATTCCGGGGACTAACGTAGATTTGGGCACTCTGGAAGGG<br>GATGGAATGAACATAGAGGGGGAGGAGTTAATGCCGTCTTTGCAAGAGGCACTGTCTTCT<br>GATATCTTAAACGATATGGAAAGCGTGTTGGCCGCCACAAAGTTGGACAAAGAATCTTTCT<br>TAACATGGTTGTAG |
| Gene Block 4 | CAACCATCTGTGAGTCCAGGGGAACCGTCTCCACCATCCATCCATCCAGGAAAATCAGACT<br>TAATAGTCCGCGTGGGCGACGAGATTAGGCTGTTATGCACTGATCCGGGCTTTGTCAAAT<br>GGACTTTTGAGATCCTGGATGAAACGAATGAGAATAAGCAGAATGAATGGATCACGGAA |

| | |
|---|---|
| | AAGGCAGAAGCCACCAACACCGGCAAATACACGTGCACCAACAAACACGGCTTAAGCAAT<br>TCCATTTATGTGTTTGTTAGAGATCCTGCCAAGCTTTTCCTTGTTGACCGCTCCTTGTATGGG<br>AAAGAAGACAACGACACGCTGGTCCGCTGTCCTCTCACAGACCCAGAAGTGACCAATTATT<br>CCCTCAAGGGGTGCCAGGGGAAGCCTCTTCCCAAGGACTTGAGGTTTATTCCTGACCCCAA<br>GGCGGGCATCATGATCAAAAGTGTGAAACGCGCCTACCATCGGCTCTGTCTGCATTGTTCT<br>GTGGACCAGGAGGGCAAGTCAGTGCTGTCGGAAAAATTCATCCTGAAAGTGAGGCCAGC<br>CTTCAAAGCTGTGCCTGTTGTGTCTGTGTCCAAAGCAAGCTATCTTCTTAGGGAAGGGGAA<br>GAATTCACAGTGACGTGCACAATAAAAGATGTGTCTAGTTCTGTGTACTCAACGTGGAAAA<br>GAGAAAACAGTCAGACTAAACTACAGGAGAAATATAATAGCTGGCATCACGGTGACTTCA<br>ATTATGAACGTCAGGCAACGTTGACTATCAGTTCAGCGAGAGTTAATGATTCTGGAGTGTT<br>CATGTGTTATGCCAATAATACTTTTGGATCAGCAAATGTCACAACAACCTTGGAAGTAGTA<br>GATAAAGGATTCATTAATATCTTCCCCATGATAAACACTACAGTATTTGTAAACGATGGAG<br>AAAATGTAGATTTGATTGTTGAATATGAAGCATTCCCCAAACCTGAACACCAGCAGTGGAT<br>CTATATGAACAGAACCTTCACTGATAAATGGGAAGATTATCCCAAGTCTGAGAATGAAAGT<br>AATATCAGATACGTAAGTGAACTTCATCTAACGAGATTAAAAGGCACCGAAGGAGGCACT<br>TACACATTCCTAGTGTCCAATTCTGACGTCAATGCTGCCATAGCATTTAATGTTTATGTGAA<br>TACAAAACCAGAAATCCTGACTTACGACAGGCTCGTGAATGGCATGCTCCAATGTGTGGCA<br>GCAGGATTCCCAGAGCCCACAATAGATTGGTATTTTTGTCCAGGAACTGAGCAGAGATGCT<br>CTGCTTCTGTACTGCCAGTGGATGTGCAGACACTAAACTCATCTGGGCCACCGTTTGGAAA<br>GCTAGTGGTTCAGAGTTCTATAGATTCTAGTGCATTCAAGCACAATGGCACGGTTGAATGT<br>AAGGCTTACAACGATGTGGGCAAGACTTCTGCCTATTTTAACTTTGCATTTAAAGGTAACA<br>ACAAAGAGCAAATCCATCCCCACACCCTGTTCACTCCT |

**Table S5.** Oligonucleotides encoding randomized peptides.

| | Sequence |
|---|---|
| Oligo 1 | 5'-GGA CAA TTA CTA TTT ACA ATT ACA ATG NNN NNN NNN NNN NNN AAA GGC AGC GGC TCC GGT CAT CAC CAC CAT CAC CAT ATG GGA ATG-3' |
| Oligo 2 | 5'-GGA CAA TTA CTA TTT ACA ATT ACA ATG NNN NNN NNN NNN NNN NNN NNN AAA GGC AGC GGC TCC GGT CAT CAC CAC CAT CAC CAT ATG GGA ATG-3' |
| Oligo 3 | 5'-GGA CAA TTA CTA TTT ACA ATT ACA ATG NNN NNN NNN NNN NNN NNN NNN NNN NNN NNN AAA GGC AGC GGC TCC GGT CAT CAC CAC CAT CAC CAT ATG GGA ATG-3' |
